# Acoustic Measures of Prosody in Right-Hemisphere Damage: A Systematic Review and Meta-Analysis

**DOI:** 10.1101/676734

**Authors:** Ethan Weed, Riccardo Fusaroli

## Abstract

The right hemisphere has often been claimed to be a locus for affective prosody, and people with right-hemisphere damage (RHD) have often been reported to show impairments in this domain. This phenomenon has been primarily investigated in terms of perception, more rarely in terms of production, and more rarely still using acoustic analysis. Our goal was to systematically review the papers reporting acoustic features of prosodic production in RHD, to identify strengths and weaknesses in this field, suggest guidelines for future research, and to support cumulative research by estimating the meta-analytic effect size of those features. We queried PubMed, PsychINFO, Web of Science, and Google Scholar, using the following combination of search terms: (prosody OR intonation OR inflection OR intensity OR pitch OR fundamental frequency OR speech rate OR voice quality) AND (RHD OR right hemisphere) AND (stroke) AND (acoustic). Standardized mean differences were extracted from all papers meeting inclusion criteria, and aggregated effect sizes were estimated using hierarchical Bayesian regression models. Sixteen papers met our inclusion criteria. We did not find strong evidence in the literature to indicate that the prosodic productions of people with RHD is substantially different from that of NBD controls, when measured in terms of acoustic features. However, the acoustic features of productions by people with RHD did differ from those of participants with NBD and LHD in some ways, notably in F0 variation and pause duration. Prosody type (emotional vs. linguistic) had very little effect. Taken together, currently available data show only a weak effect of RHD on prosody production. However, more accurate analyses are hindered by small sample sizes, lack of detail on lesion location, and divergent measuring techniques. Cumulative open science practices are recommended to overcome these issues.

## 1. Introduction

Speakers modulate their tone of voice to add important nuance to their words. This could range from changes of emphasis which carry lexical distinctions (e.g. a *hot*dog vs. a hot *dog*) or syntactic distinctions (e.g. a terminal upward inflection to indicate a question) to an overall colouring of tone indicating emotion or a particular attitude toward the uttered words (e.g. anger, sarcasm, amusement, etc.). Since the 19th century, clinicians have recognized that the loss of this ability can have important implications for communication (Hughlings Jackson, 1879), and since the early 20th century, researchers have attempted to identify and classify the various types of disordered prosody (Monrad-Krohn, 1957; Monrad-Krohn, 1947). In the later 20^th^ century, perhaps in conjunction with a general upsurge of interest in communication deficits following damage to the brain’s right hemisphere (Gardner, 1994; Myers, 1999; Tompkins, 1995; Winner, Brownell, Happé, Blum, & Pincus, 1998), attention turned to the right hemisphere as a potential neural locus of at least some aspects of prosody, with Ross in particular (Gorelick & Ross, 1987; Ross, Harney, & Purdy, 1981; Ross & Mesulam, 1979; Ross, Thompson, & Yenkosky, 1997) describing several patients whose prosody became “monotone” following damage to the right hemisphere, and proposing that affective prosody might be a dominant language function of the right hemisphere. One particular case described by Ross and Mesulam (Ross & Mesulam, 1979) was that of a school teacher who, following a right hemisphere stroke, had a “complete inability to express emotion through speech and action,” and who spoke with an “asthenic, unmodulated, monotonous voice that was devoid of inflections and coloring” (pg. 144). This individual compensated by adding “and I mean it” to the end of sentences to indicate anger (Ross & Mesulam, 1979).

In subsequent years, the right hemisphere has come to be widely associated with prosody, and with emotional or affective prosody in particular (Guranski & Podemski, 2015; Patel et al., 2018; Leung, Purdy, Tippett, & Leão, 2017). Indeed, Guranski and Podemski (2015) write that the right hemisphere is “[…] mostly responsible for speech prosody and its emotional aspects” (pg. 113). The focus shifted to the perception of emotional prosody, rather than production (Patel et al., 2018), and was reinforced by neuroimaging studies on healthy participants processing emotional prosody, e.g. (Baumgaertner, Hartwigsen, & Roman Siebner, 2013). Even the occasional studies including a production component did not typically consider the acoustic properties of production, but rather measured prosody using hearer judgments, e.g (Brådvik et al., 1991).

Despite the reported case studies, and the general acceptance that the right hemisphere has a special role to play in prosody processing, no comprehensive systematic literature review or meta-analysis yet exists on the role of the right hemisphere in prosodic processing in general, or even in production specifically. The present study takes a first step in this direction by carrying out the first systematic review and meta-analysis of studies using acoustic measures of prosodic production in people with right hemisphere damage (RHD). The goal is to identify strength and weaknesses in this field and thereby guidelines for future research, as well as to support cumulative research by estimating meta-analytic effect size of those features.

In studies using hearer judgments to assess prosodic production, patients are asked to e.g. produce either compound nouns or noun phrases (hotdog vs. hot dog). Raters then listen to these productions, and classify them as either a compound noun or noun phrase (e.g. Balan & Gandour, 1999). Similar procedures are used for emotional prosody, e.g. in Gandour, Larsen, Dechongkit, Ponglorpisit, & Khunadorn (1995), in which raters circled happy, sad, or neutral smileys to classify patient productions. Studies using acoustic measures, in contrast, extract features directly from the recorded speech signal. These physical characteristics of the speech signal are experienced by the listener as elements of prosody. Acoustic features include fundamental frequency (pitch), intensity (loudness), and duration measures (pauses and rhythm).

We have chosen to focus on acoustic measures because we believe these to be the most promising means currently available for assessing prosody in speech production. We are not alone in this: previous reviews on perceptually assessed prosodic patterns in Autism Spectrum Disorder and Schizophrenia have suggested moving toward acoustic analyses, in order to provide more objective and mechanistic descriptions of the prosodic atypicalities in these disorders (Fusaroli, Lambrechts, Bang, Bowler, & Gaigg, 2017; Hoekert, Kahn, Pijnenborg, & Aleman, 2007; McCann & Peppé, 2003; Parola, Simonsen, Bliksted, & Fusaroli, 2019). This is not to suggest that studies based on hearer judgements are not valid or useful. Indeed, we believe that hearer judgments do provide an important source of information, as they allow us to quantify the subjective impression that a person’s speech makes on a listener. Insofar as atypical speech patterns can be considered an impairment at all, it is only to the extent to which they hinder the hearer’s ability to quickly and accurately comprehend the intended message of the speaker, build rapport with the speaker, and coordinate the flow of the conversation. These can be better assessed through hearer judgments, or by other means which reflect the effect of the acoustic cues on the hearer or the conversation. However, hearer judgements are also limited in several respects:

1. The data generated by hearer judgements tends to be categorical, or at the very least ordinal. Acoustic measures are continuous, allowing for more sensitive analysis methods (Bürkner & Vuorre, 2018).
2. Hearer judgments synthesize potentially many separate acoustic cues into a single measure. As an example, many different acoustic features may contribute to an overall “monotone” quality of speech, but the specific features may differ from speaker to speaker (Forbes-Riley & Litman, 2004; Liscombe, Venditti, & Hirschberg, 2003).
3. Acoustic analysis of speech offers many more potential measures than ratings-based judgements. For example, a commonly used suite (COVAREP) provides 82 out-of-the-box acoustic measures, which have been repeatedly demonstrated to be useful for describing voice patterns (Degottex, Kane, Drugman, Raitio, & Scherer, 2014). These measures – when coupled with conservative statistics to avoid overfitting - may help better characterize subtle differences in speech patterns.
4. Hearer judgments do not scale well. While it is possible to ask raters to judge the speech of a small sample of participants, as data-sets become larger and more openly shared (e.g. at https://rhd.talkbank.org (Macwhinney, Fromm, Forbes, & Holland, 2011)), it is increasingly important to develop replicable methods for extracting measures from many sound files in a uniform fashion. Using standardized, freely-available tools and providing analysis code allows researchers to more easily build upon each other’s work.

Although we have chosen to focus on studies using acoustic measures to assess prosody production, there is increasing interest in combining hearer judgments and acoustic measures to identify acoustic correlates of perceived prosody (e.g. Jiam, Caldwell, Deroche, Chatterjee, & Limb, 2017; Moriarty, Vigeant, Liu, Gilmore, & Cole, 2018; Stoop et al., 2018), and we believe this method holds promise for work on prosody in clinical populations as well.

The aim of the current meta-analysis is to assess whether, and to what degree, acoustic features of the speech of people with RHD differ from those same features in the speech of non-brain-damaged (NBD) controls, by drawing on currently available results in the literature.

## 2. Methods

### 2.1 Literature search

To obtain the most comprehensive database of peer-reviewed results possible, we conducted a systematic literature search. We queried PubMed, PsychINFO, Web of Science, and Google Scholar, using the following combination of search terms: (prosody OR intonation OR inflection OR intensity OR pitch OR fundamental frequency OR speech rate OR voice quality) AND (RHD OR right hemisphere) AND (stroke) AND (acoustic), with no restrictions on date of publication. Further papers were identified by searching the references in the papers identified by the initial search. Our search resulted in a total of 49 papers.

### 2.2 Inclusion criteria

Our initial list of 49 papers was then screened, and we included only papers meeting the following criteria:

1. The study quantified acoustic measures of vocal production of participants with RHD
2. The study reported original research (not a review)
3. The study included data from at least two participants
4. The study included a control group

Of the initial 49 papers, 8 were case studies with data from only one individual, 6 did not report sufficient data in either tables or plots for inclusion in a meta-analysis, 2 papers did not report data from an acoustic analysis, 2 papers lacked a control group, 11 papers did not measure vocal production, 2 did not include participants with RHD, 1 did not report original data (review article), and 1 only included RHD participants who were actively selected to differ from NBD controls. Although this final paper otherwise met our inclusion criteria, we felt that actively choosing patients who differed from the control group on the object of study introduced unnecessary bias, and we did not include it. We note in passing that the effect sizes for this paper (which measured variation in F0) were somewhat larger (Hedges’s *g* = 1.3 – 1.8) than many of the included papers, although not unlike the effect size in a paper with the same first author which was included (Ross & Monnot, 2008). After these 33 papers were excluded, 16 papers remained for analysis. Full details on these papers and the features measured can be found in the data-table archived at: https://osf.io/2g8fr/?view_only=81ffc526d83c4389b5b4fc073c2c922a.

### 2.3 Data extraction

The elementary acoustic features usually argued to support prosody are fundamental frequency (F0), intensity, and duration (Fusaroli et al., 2017; Jiam et al., 2017; Peppé, 2009). These acoustic features are interpreted by the hearer as pitch, loudness, and rhythm or timing. Reported data on any of these features were extracted for our analysis. In many cases, these data could be easily extracted from tables published in the articles. In some cases, sufficient data were not available in table format, but could be digitally extracted from published plots using the application WebPlotDigitzer (Rohatgi, 2014). In one case, a printing error had resulted in published data tables which were clearly incorrect; in this case, the authors were contacted and kindly provided the correct data. Because lesion location was either unreported or was reported as a mix of varied cortical and subcortical damage, there was insufficient data to include lesion location as a predictor in our models. When papers also reported results for participants with left-hemisphere damage (LHD) these were also extracted.

### 2.4 Meta-analysis

Meta-analyses were performed following well-established random- and mixed-effects procedures detailed in Doebler & Holling (2015); Field & Gillett (2010); Quintana (2015); and Viechtbauer (2010) complemented – in the inference of the effects - by a Bayesian framework (Williams, Rast, & Bürkner, 2018). To estimate the differences between individuals with RHD and non-brain-damaged (NBD) controls, we extracted the standardized mean difference (Hedges’ *g* (Borenstein, Hedges, Higgins, & Rothstein, 2011; Cohen, 1988)). When papers reported data from more than one measure of the same variable, each measure was entered separately into the calculation. We refer to these multiple measurements of the same variable as “studies”, while the publications they are reported in we refer to as either “articles” or “papers”.

The standardized mean differences were analysed using 2-level hierarchical Bayesian regression models to estimate the pooled effect sizes and corresponding credible (i.e., Bayesian confidence) intervals. This multilevel structure allowed us to explicitly model the heterogeneity (or σ^2^) in the results of the studies analysed. By including a random effect by study, we both accounted for repeated measures and assumed that the variability in experimental design, acoustic analyses, and population samples might generate heterogeneous findings, and allowed the model to estimate such heterogeneity (Hedges, Tipton, & Johnson, 2010).

We then measured and tested for heterogeneity between the studies using the Cochran’s Q statistic (Cochran, 1954). Q statistics reveal how much of the overall variance can be attributed to true between-study variance. Priors for the Bayesian analyses were chosen to be only weakly informative, so that their influence on the meta-analytic estimates were relatively small: a normal distribution centred at 0 (no effect), with a standard deviation of 0.5 for the overall effect, and a positive truncated normal distribution centred at 0, with a standard deviation of 0.5 for the heterogeneity of effects (standard deviation of random effects). We report 95% credible intervals (CIs), evidence ratios and credibility scores. CIs are the intervals within which there is a 95% probability that the true value of the parameter (e.g. effect size) is contained, given the assumptions of the model. In the present analysis, the evidence ratio quantifies evidence in favour of the effect of diagnosis or of acoustic feature (e.g. longer pauses in RHD compared to NBD) and provides the ratio of likelihood between this effect and the alternatives (e.g. same length or shorter pauses in RHD). An evidence ratio equal to 3 indicates the hypothesis is 3 times more likely than the alternative. A common interpretation of evidence ratios, proposed by Wetzels and Wagenmakers (2012) and based on work by Jeffreys (1998) and Kass and Raftery (1995), is as follows: 1–3 = anecdotal; 3–10 = substantial; 10–30 = strong).

A credibility score indicates the percentage of posterior estimates falling above 0. Because Bayesian methods are less commonly used and understood, we also report p-values in order to reach a broader audience. Note that the p-values are calculated on the same 2-level hierarchical model as the Bayesian inference, with the difference that p-value statistics rely on the equivalent of completely flat priors.

The presence of influential studies is estimated using Cook’s distances, that is, per each study we calculate the Mahalanobis distances (Mahalanobis, 1936) between the entire set of predicted values and the set of all but the current study. Studies generating a Cook distance above 1 are considered influential and the data are subsequently re-analysed excluding all influential studies (Cook & Weisberg, 1982; Viechtbauer, 2010).

To assess the potential role of prosody type (emotional vs. linguistic prosody) in explaining the patterns observed, we built a second 2-level Bayesian model including prosody type as predictor of difference in vocal patterns. We then performed a Leave-One-Out-Information-Criterion (loo) based model comparison (Vehtari, Gelman, & Gabry, 2017) with the model not including task. We report loo stacking weights (Yao, Vehtari, Simpson, & Gelman, 2018) in favour of the model, indicating the probability that the model including the variable *prosody type* is better than baseline.

Publication bias was assessed by both visual inspection of asymmetries in the funnel plots, and Egger’s regression test (Egger, Smith, Schneider, & Minder, 1997). All calculations were carried out using the metafor (Viechtbauer, 2010) and brms (Bürkner, 2017a; Bürkner, 2017b) packages for R (R Core Team, 2018).

Meta-analyses were conducted comparing both participants with RHD to NBD controls and, when possible, to participants with LHD. This latter comparison helps identify differences between RHD and NBD participants which might simply be the result of any sort of brain damage, irrespective of hemisphere.

## 3. Results

Not all of the extracted features were reported in a sufficient number of papers to make meta-analysis possible. Meta-analytic comparison between RHD and NBD participants was possible for the features F0 variability, Pause Duration, Syllable Duration, and Intensity Duration. These same comparisons were also possible between RHD and LHD participants, with the exception of Syllable Duration. The main results of the analyses are summarised in Table 1. Below, we look at each of these results in more detail.

**Table 1:**
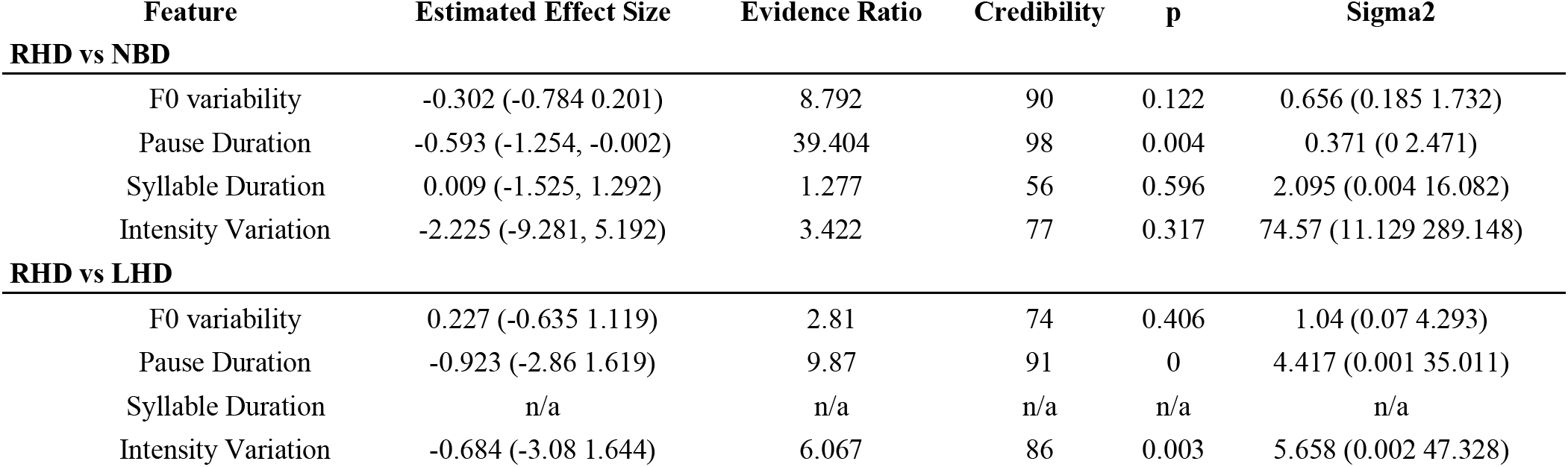
Meta-analytic results for comparisons of RHD vs. NBD and (where possible) RHD vs. LHD.

### 3.1 Variability in Fundamental Frequency (F0)

The most commonly-studied aspect of prosody in the papers we included in our meta-analysis was pitch, in particular pitch variation. As only two studies (Behrens, 1988; Pell, 1999a) reported mean F0, and only one study reported measures of pitch contour (Behrens, 1989), there were not sufficient data to meaningfully run a meta-analysis on measures of mean pitch or pitch contour. We have made these data available, however, in the online data-table: https://osf.io/2g8fr/?view_only=81ffc526d83c4389b5b4fc073c2c922a.

The meta-analysis of variation in F0 included 29 studies (13 articles) for a total of 167 participants with RHD and 187 comparison participants (Figure 1). Hierarchical Bayesian meta-analysis revealed an overall estimated difference (Hedges’ g) in F0 Variability of −0.302 (95% CI = - 0.784, 0.201), p = 0.122, evidence ratio = 8.792, credibility = 90%) with an overall variance between studies (sigma squared) of 0.656 (95% CI = 0.185, 1.732). The variance in effects between studies could not be reduced to random sample variability between studies (Q-stats: 110.478, p = 0). No study was found to be influential. The data did not reveal any likely publication bias (Kendall’s tau = 0.069, p = 0.616). Adding prosody type did not improve the model (Stacking weight: 0%).

**Figure 1:**
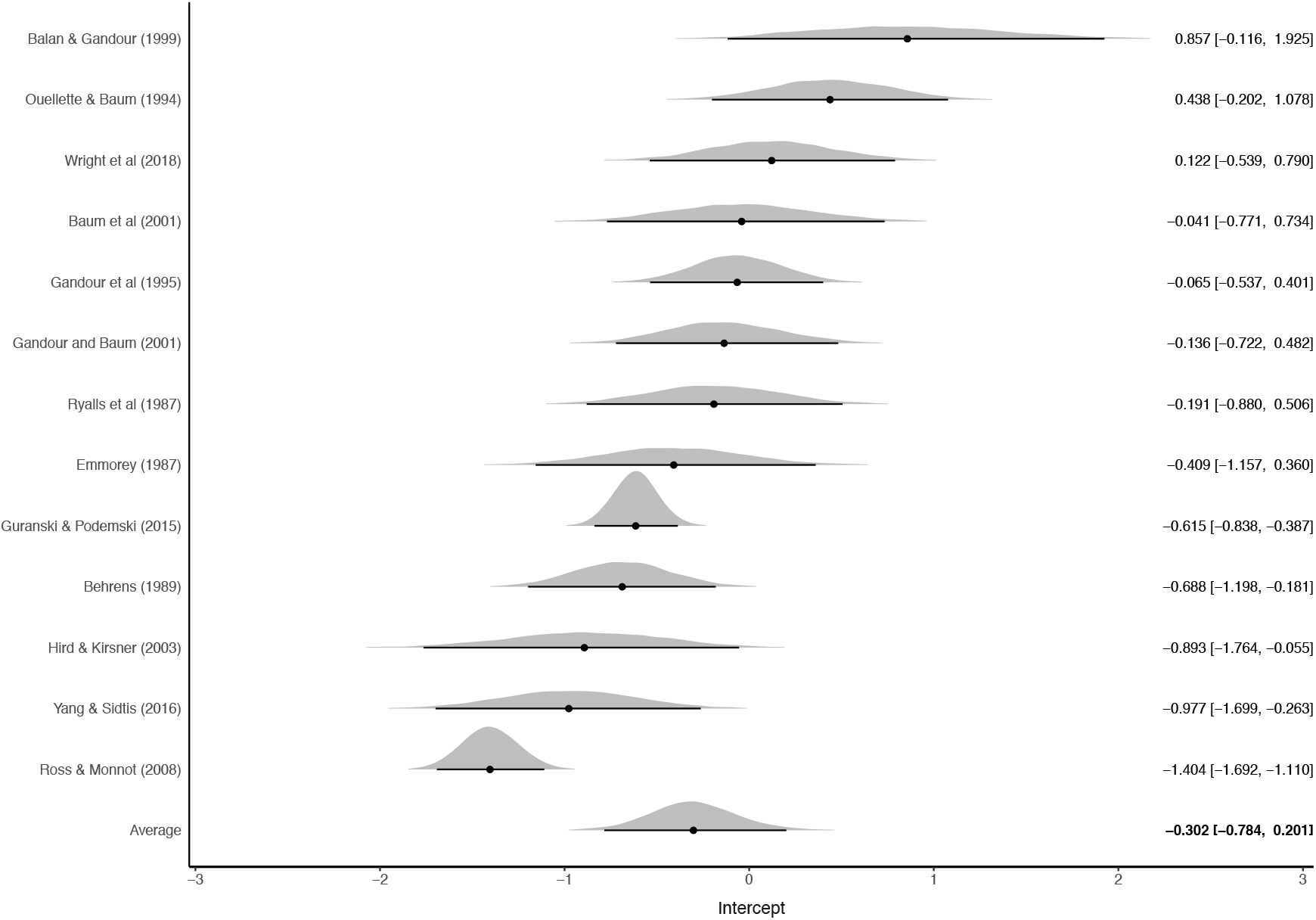
Estimated effect sizes for variability of F0. Shaded areas indicate the posterior probability density of each estimate. Numbers indicate estimated mean difference (Hedge’s g) and upper and lower 95% credibility intervals.

### 3.2 Pause Duration

The meta-analysis of mean pause duration included 7 studies (4 articles) for a total of 28 participants with RHD and 35 comparison participants (Figure 2). Hierarchical Bayesian meta-analysis revealed an overall estimated difference (Hedges’ g) in Pause Duration of −0.593 (95% CI = −1.254, −0.002), p = 0.004, evidence ratio = 39.404, credibility = 98%) with an overall variance between studies (sigma squared) of 0.371 (95% CI = 0, 2.471). The variance in effects between studies could be reduced to random sample variability between studies (Q-stats: 0.898, p = 0.989). No study was found to be influential. The data did not reveal any likely publication bias (Kendall’s tau = −0.429, p = 0.239).

**Figure 2:**
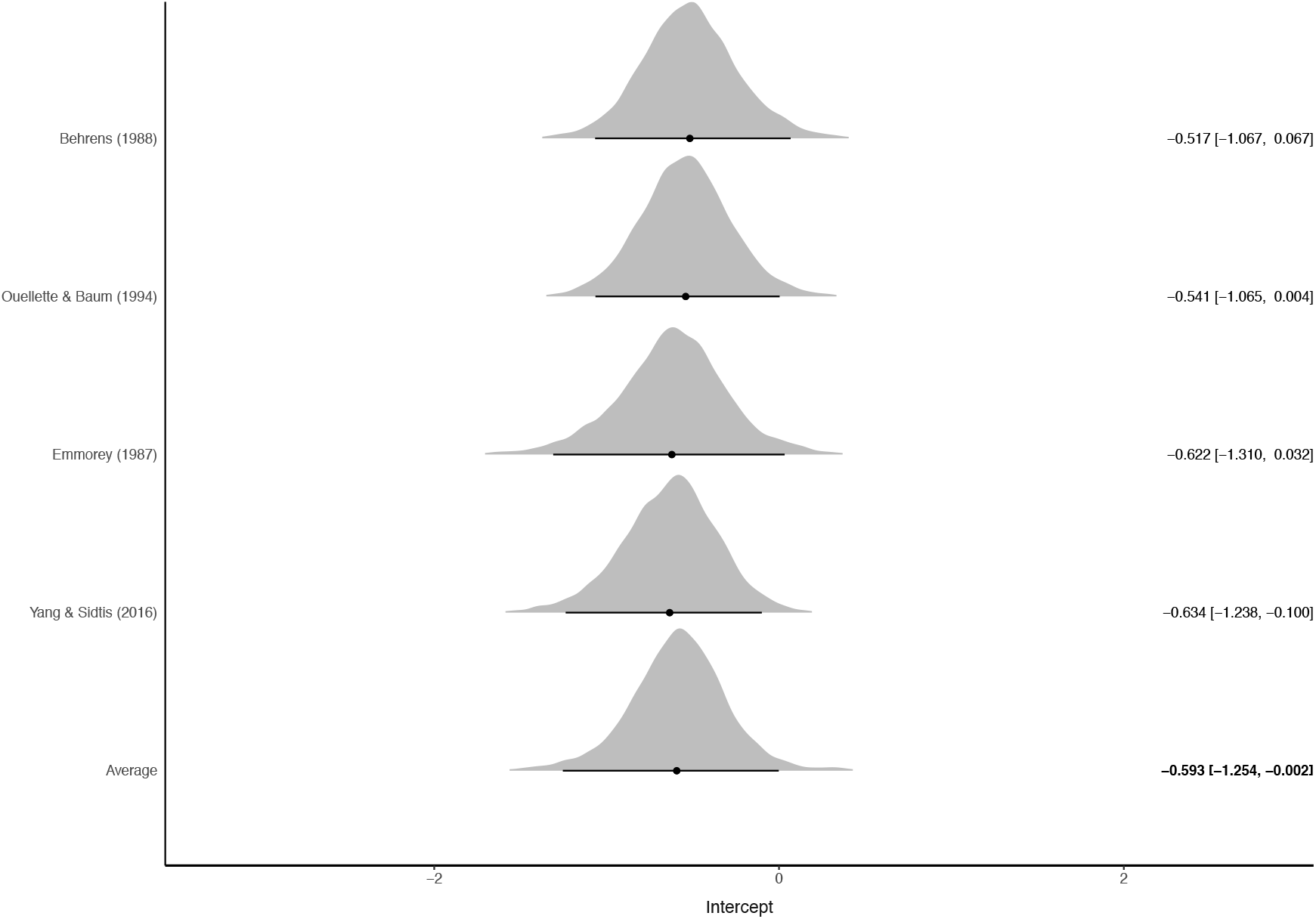
Estimated effect sizes for Pause Duration. Shaded areas indicate the posterior probability density of each estimate. Numbers indicate estimated mean difference (Hedge’s g) and upper and lower 95% credibility intervals.

### 3.3 Syllable Duration

The meta-analysis included 13 studies (4 articles) for a total of 32 participants with RHD and 46 comparison participants (Figure 3). Hierarchical Bayesian meta-analysis revealed an overall estimated difference (Hedges’ g) in Syllable Duration of 0.009 (95% CI = −1.525, 1.292, p = 0.596, evidence ratio = 1.277, credibility = 56%) with an overall variance between studies (sigma squared) of 2.095 (95% CI = 0.004, 16.082). The variance in effects between studies could be reduced to random sample variability between studies (Q-stats: 11.697, p = 0.47). No study was found to be influential. The data revealed a likely publication bias (Kendall’s tau = 0.692, p = 0.001). Adding prosody type did not improve the model (Stacking weight: 0%).

**Figure 3:**
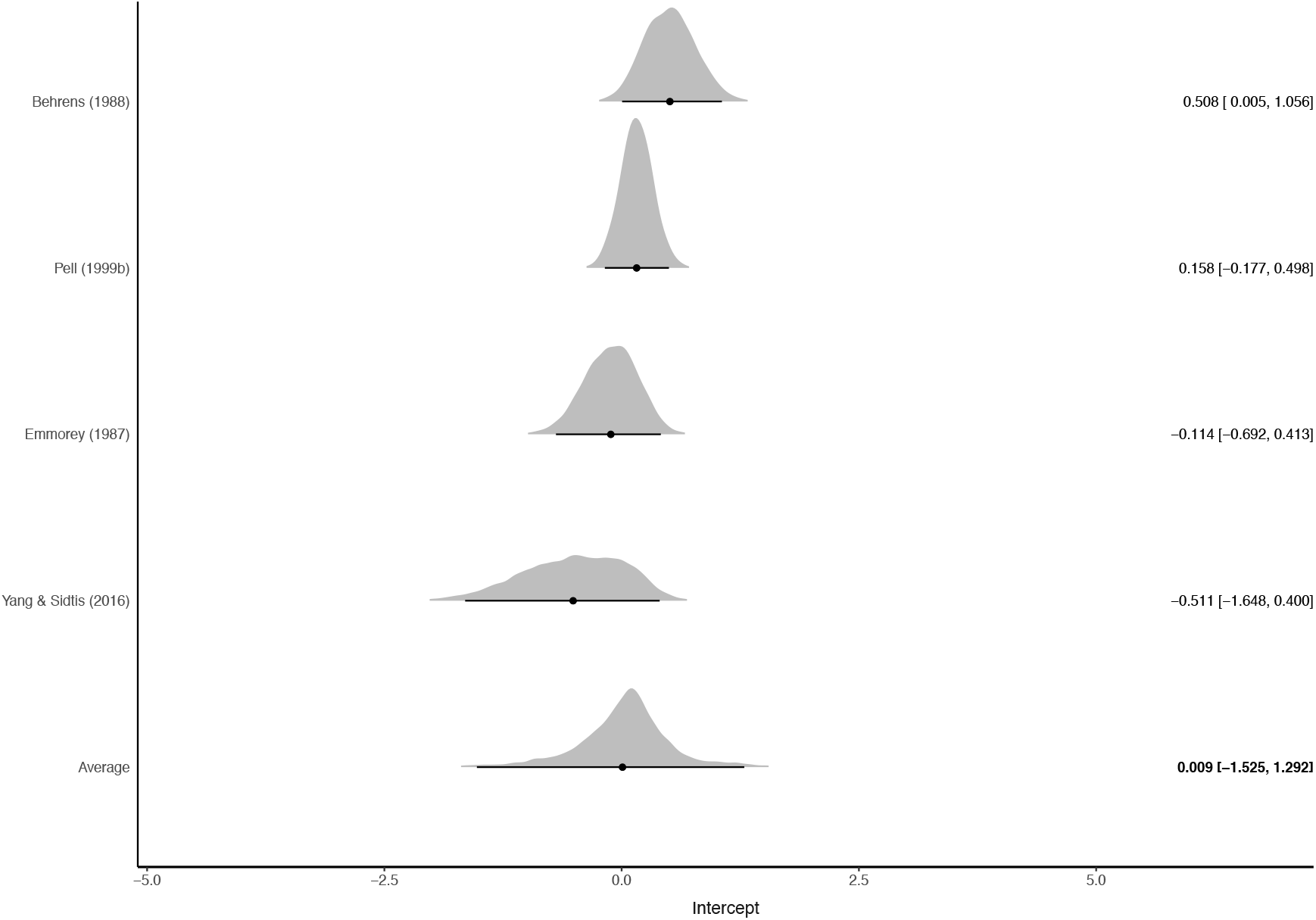
Estimated effect sizes for Syllable Duration. Shaded areas indicate the posterior probability density of each estimate. Numbers indicate estimated mean difference (Hedge’s g) and upper and lower 95% credibility intervals.

### 3.4 Variability in intensity

The meta-analysis included 9 studies (5 articles) for a total of 46 participants with RHD and 43 comparison participants (Figure 4). Hierarchical Bayesian meta-analysis revealed an overall estimated difference (Hedges’ g) in Intensity Variation of −2.225 (95% CI = −9.281, 5.192, p = 0.317, evidence ratio = 3.422, credibility = 77%) with an overall variance between studies (sigma squared) of 74.57 (95% CI = 11.129, 289.148). The variance in effects between studies could not be reduced to random sample variability between studies (Q-stats: 39.182, p = 0). No study was found to be influential. The data did not reveal any likely publication bias (Kendall’s tau = 0.111, p = 0.761). The data did not reveal any likely publication bias (Kendall’s tau = 0.111, p = 0.761). Adding prosody type to the model credibly improved it (Stacking weight: 100%). Emotional prosody showed an effect of 0.22 (95% CI = −17.701, 18.766), while linguistic prosody showed an effect of −3.154 (95% CI = −12.297, 6.257), indicating that participants with RHD show slightly more variability in intensity than controls when producing emotional prosody, but substantially less than controls when producing linguistic prosody. Note that this is due to only one study (Hird & Kirsner, 2003) reporting very large effects and should therefore interpreted with much caution.

**Figure 4:**
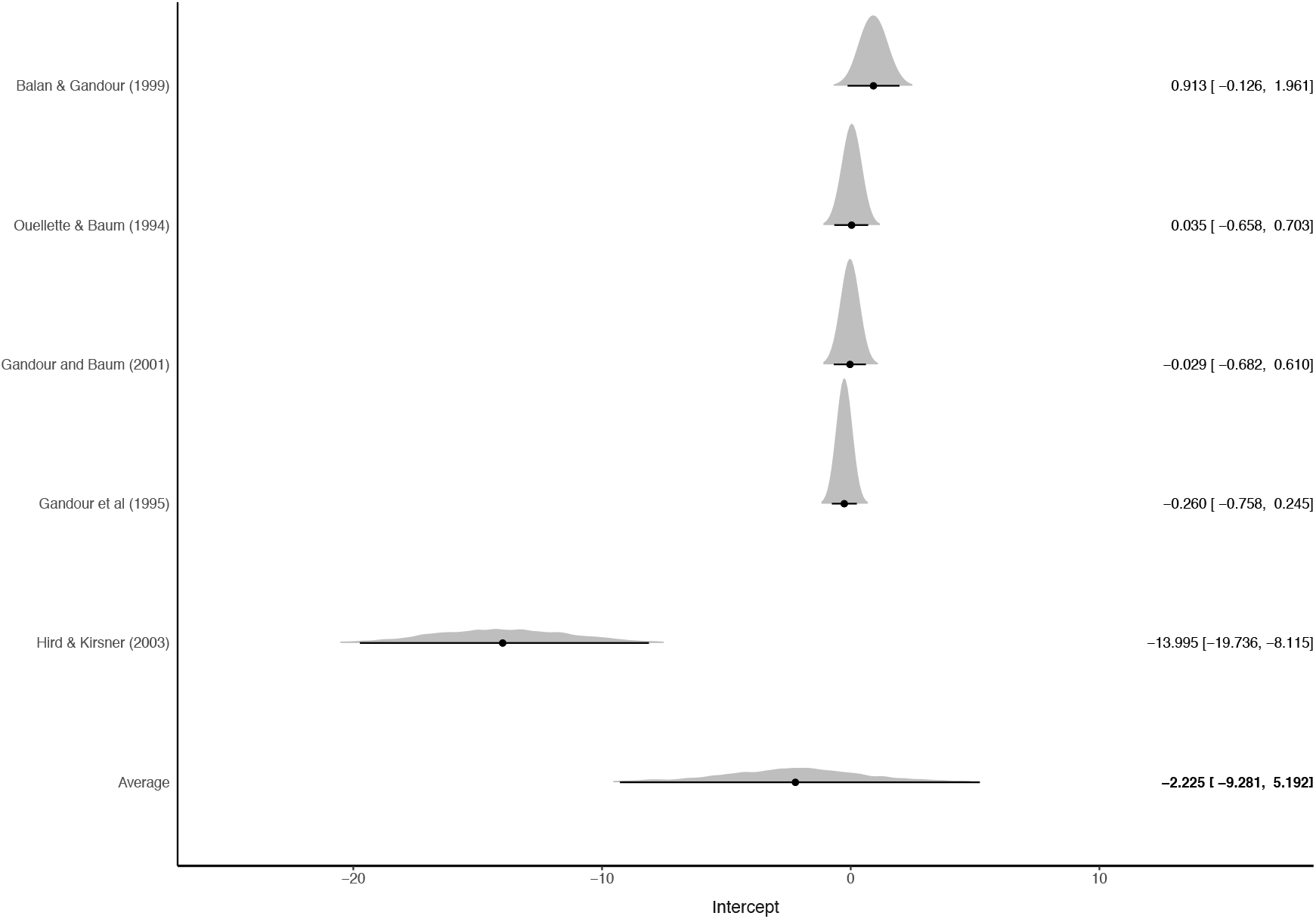
Estimated effect sizes for variability for variability of intensity. Shaded areas indicate the posterior probability density of each estimate. Numbers indicate estimated mean difference (Hedge’s g) and upper and lower 95% credibility intervals.

## 4. Discussion

The purpose of this meta-analysis was to evaluate to what degree prosody, as measured by acoustic analysis, is related to right-hemisphere damage. We extracted data from all available articles comparing data on acoustic measures of pitch, rhythm, and intensity from people with RHD with that of NBD controls. Where possible, we have also included data from people with LHD. Taken together, the data from acoustic studies reveal some, but not substantial, differences in prosody production in people with RHD when compared with NBD controls. Below, we discuss the findings and their generalisability, and make recommendations for future studies.

The most commonly measured feature was pitch variance. We found moderate evidence for reduced variation in pitch in people with RHD, and this was not moderated by prosody type (linguistic vs emotional). Because the result did not appear to be driven by any particularly influential studies, the lack of evidence of publication bias, and the highly credible estimate, we suggest that this result is reasonably sound: pitch variation is reduced in people with RHD, although the effect is small. While this does support the idea that people with RHD tend to speak in a monotone, the meta-analysis also indicated that variation in pitch was essentially the same for people with RHD and LHD (in fact, variation was slightly higher for people with RHD, see Table 1), suggesting that the effect may be due to issues surrounding brain damage in general, and not specific to damage to the RH.

The strongest and most convincing effect was that of pause duration. Although only four articles measured pause duration, the results were surprisingly consistent. It is important to note that pause duration was very narrowly defined in these articles, and was not entirely consistent. The older articles (Behrens, 1988; Emmorey, 1987; Ouellette & Baum, 1994) measured pause duration in the context of noun-noun compounds and noun phrases, and defined pause duration as the time between the last glottal pulse of the first syllable’s vowel and onset of the second syllable’s vowel. Yang and Sidtis (Yang & Van Lancker Sidtis, 2016), in contrast, defined pause duration as the percentage of pause duration in the whole sentence in a task involving the production of idiomatic vs. literal sentences. No articles reported measures of pause duration in the context of emotional prosody. Interestingly, of the papers included in the meta-analysis, only Behrens (Behrens, 1988) concludes that participants with RHD use pause differently than that of controls, and Behrens’ conclusion is challenged by Ouellette and Baum (Ouellette & Baum, 1994)). The results of the meta-analysis suggest that, at least within a narrowly-defined context, pause duration is indeed affected in RHD.

Not only did the participants with RHD produce shorter pauses than the NBD participants, their pauses were also shorter than LHD participants (Table 1). Although the evidence for this finding is weaker than that for the comparison of RHD to NBD participants, it is an important finding, as it indicates that the effect seen in participants with RHD is not simply due to brain damage in general, but may be particularly pronounced following damage to the right hemisphere. This compression of syllables has been earlier described as a “fused” pattern by Kent and Rosenbek (1982), who identify the pattern as being typical of both patients with RHD and patients with Parkinson’s disease, and describe it as a form of dysarthria.

We found no evidence for an effect of RHD on vowel or syllable duration when compared with NBD controls, with an estimated effect size of essentially zero. This finding should be considered in light of the narrow definition used in this analysis. Of the 6 articles that measured some aspect of speech duration, several different types of measurement were employed. These included measures of utterance length (Gandour et al., 1995; Guranski & Podemski, 2015), variability of syllable duration (Yang & Van Lancker Sidtis, 2016), and mean vowel or syllable duration (Behrens, 1988; Emmorey, 1987; Pell, 1999b; Yang & Van Lancker Sidtis, 2016). We considered these measures too diverse to be meaningfully combined, and of these only vowel or syllable duration was represented by enough papers to conduct a meta-analysis. It may well be the case that some other measure of duration, e.g. utterance duration or articulation time, would reveal an effect.

Variability of intensity was the only measure of intensity with sufficient data to conduct a meta-analysis. We found no evidence of RHD on variability of intensity when compared with NBD controls. Interestingly, participants with RHD did show substantially less variation in intensity when compared with participants with LHD. At the same time, reported intensity data should always be treated with caution, as these measurements can be affected by a variety of factors including the choice of microphone, and the distance of the microphone from the speaker. An obvious outlier in the intensity data is the study by Hird and Kirsner (2003). This study measured the change in intensity between breath groups during informal conversation, unlike e.g. Balan and Gandour (1999) who examined intensity changes between syllables in a phonemic stress paradigm, with elicited production of noun-noun compounds.

Although many authors distinguish between linguistic prosody (e.g. using intonation to distinguish between noun-noun compounds and noun phrases) and emotional or affective prosody, this distinction did not play a major role in the meta-analysis. Only one model, intensity, was credibly improved by adding prosody type, but credibility intervals were still so large that these results were difficult to interpret with any confidence. Furthermore, measures of intensity can be difficult to compare across studies, as this feature can be greatly affected by the methods and materials used in data collection. Although RHD has been at times associated with changes in emotional rather than linguistic prosody (for a brief review see (Kotz et al., 2003)), our strongest finding was from a feature only measured in studies investigating linguistic prosody. Whether this feature would also be affected in the context of emotional prosody is unknown.

When evaluating these results, it is important to keep a few points in mind. First, the studies included in the present meta-analysis are group-level analyses, and do not report pre- vs. post-trauma data within individuals. Second, because of the lack of available information on lesion location, we have considered the hemisphere as the unit of analysis. This reflects the literature, which has not always been able to make more fine-grained distinctions, although these would undoubtedly paint a more accurate picture. Finally, although the papers we have included in our analysis report unidimensional measures of prosody, it should be noted that different acoustic measures can be used to similar effect. For example, emotions can be indicated by modulating pitch, intensity, pauses, or all three. By systematically reviewing the literature, we were able to identify these cautions, which should be kept in mind for future studies (e.g. including more precise reports of lesion location). Further, despite these cautions, it is important to evaluate the existing data within a meta-analytic framework, not only to estimate baseline effect sizes, but also to allow the selection of informed priors for future analyses, thus cumulatively incrementing our knowledge of prosody in RHD (Cumming, 2014; Gelman, Jakulin, Pittau, & Su, 2008; Williams, Rast, & Bürkner, 2018).

A common finding for all features was that effect sizes were relatively small. Although variation in intensity had a large estimated effect size of −2.225, this estimate seems unlikely to be very reliable. The most robust estimated effect size, pause duration, was at best of medium size (Hedges g: −0.593). Another common finding was that sample sizes were quite small, with a mean N of 12.16 (SD: 10.05). These small samples lead to poor estimates, both within each study, and when attempting to combine studies. Figure 1, showing estimated effect sizes for variation in F0 illustrates this clearly. In this figure, the two studies with the largest sample sizes (Guranski & Podemski (2015) with an N of 46, and Ross & Monnot (2008) with an N of 21) have the most normal-shaped estimated posterior distributions, whereas a study like Balan and Gandour (1999), with an N of 8 has a posterior distribution whose probability mass is spread out over a wide area. Although other factors are also at play, these small sample sizes make it difficult to make a reliable estimate of the true effect size.

The majority of the studies analyzed include 10 or fewer patients, plausibly due to the difficulty in accessing clinical populations. However, with such small sample size one would need a standardized mean difference (effect size) of 1.7 or more to reach a 95% power (calculations relying on G*Power (Erdfelder, Faul, & Buchner, 1996)) at which effect size estimates are reliable. This is clearly not the case for the acoustic features we have analyzed.

While including as varied a sample as possible is an unavoidable concern, there are strategies to reduce the sample size needed. For instance, one could employ repeated measures, that is, collecting repeated voice samples over time. Using even just 3 repeated measures per participant reduces the required underlying effect size from 1.7 to 1 (assuming the participants are still representative of the full population). Repeated measures are also very useful to better understand the reliability of the acoustic patterns over re-testing and potentially across different contexts. In particular, we have seen that linguistic and emotional prosody might not be as different as expected, but without a controlled within-subject contrast it is difficult to assess whether this is due to other confounds in the sample and study design.

At the same time, we recognize that it can be difficult to achieve large sample sizes, particularly when working with clinical populations. Collecting data from these participants involves not only finding participants who meet the inclusion criteria, but also requires making demands on the sometimes already taxed time and attention of the participants and their families or clinical staff. It is therefore all the more important that when data on communication disorders are collected, that they be shared so that other researchers can build on them, and so that the impact that each participant can have by donating their time to participation in research can be extended (MacWhinney, Fromm, Rose, & Bernstein Ratner, 2018). This can be e.g. by contributing data to resources such as RHDBank, a part of the TalkBank system (Macwhinney et al., 2011). Similar prosody and voice-quality datasets can already be found for e.g. Autism Spectrum Disorder (Schuller et al., 2013), Parkinson’s Disease (Tsanas, Little, McSharry, & Ramig, 2010), and Depression (Cummins et al., 2015). Although data protection legislation may make this type of data sharing more difficult, it is important that researchers adapt their practices and find ways to responsibly and legally share data, to the extent possible.

A related issue is the importance of making analysis code publicly available. The researchers whose work we have drawn on in the present meta-analysis have used a variety of methods to extract values for acoustic features, and to calculate statistics based on those values. By using publicly-available, open source tools such as COVAREP (Degottex, Kane, Drugman, Raitio, & Scherer, 2014) and by making in-house extraction algorithms or analysis code available, researchers can leave a trail for future researchers to follow and systematically build on their findings. An additional advantage of automated tools is the large number of acoustic features which can be extracted. While earlier researchers needed to manually extract e.g. pitch curves from spectrograms, automated tools can rapidly extract a wide variety of acoustic features from many sound files.

Data and analysis code from the present meta-analysis are both available at: https://osf.io/2g8fr/?view_only=81ffc526d83c4389b5b4fc073c2c922a. We make these available not only in the interest of transparency, but also as a tool for future research. Like all meta-analyses, our results reflect choices about which data to include and to exclude. We have attempted to be as clear as possible about our choices, but recognize that other authors might choose to analyse the data differently. As an example, if a future researcher is interested in studying variation in intensity in people with RHD, not only can they refer to this article for a meta-analytic estimate of effect size, they can also use the provided data and code to re-run the analysis themselves, and could choose, for instance, to exclude the study by Hird and Kirsner (2003), on the grounds that it reports data from free conversation, while they are interested in noun-noun compounds.

Finally, where possible, it will be important for future researchers to collect as accurate localisation data as possible. In the papers we have included in the meta-analysis, patients are often simply classified as having suffered a right CVA, without further specification. While it may be possible to make general claims about right- vs. left-hemisphere processes, it is also likely that the data we have analysed here have been muddied by including patients with very different lesion locations and sizes. Some patients may have anterior damage, some posterior, some may have cerebellar or subcortical damage, while others may have fairly restricted cortical lesions. These differences undoubtedly have an impact on the production of prosody, and future attempts to understand the localization of systems supporting prosody production will need more fine-grained information on lesion location.

In sum, our recommendations for future research are:

1. Counteract small sample sizes by making data publicly available in repositories, with the necessary cautions due to anonymization and data protection, or at the least share extracted acoustic features.
2. Make feature-extraction and analysis code available for reproducibility and easier comparison of data. Use standardized acoustic extraction, at least as a baseline to facilitate comparison with previous results.
3. When possible, include precise information on lesion location.

## 5. Conclusions

In this meta-analysis, we employed hierarchical Bayesian meta-analysis to evaluate the effect of right-hemisphere damage on prosodic production, as measured by acoustic analysis. Taken together, the results did not strongly support the claim that prosodic production is markedly different in people with RHD when compared with NBD controls. We found some evidence for a reduction in variability of F0 in RHD, and substantial evidence (albeit from a small number of studies) for a decrease in pause duration in RHD. This decrease in pause duration was primarily based on studies comparing the linguistic prosody distinguishing noun-noun compounds from noun phrases. Although we found studies spanning nearly 30 years measuring acoustic features of prosodic production in RHD, there is still relatively little data available to draw any strong conclusions. This situation could be aided by increased attention to algorithmic feature extraction and open data sharing.

## References

Balan, A., & Gandour, J. (1999). Effect of sentence length on the production of linguistic stress by left-and right-hemisphere-damaged patients. Brain and language, 67(2), 73–94.

Baum, S. R., Marc D. Pell, Carol. (2001). Using prosody to resolve temporary syntactic ambiguities in speech production: acoustic data on brain-damaged speakers. Clin Linguist Phon, 15(6), 441–456.

Baumgaertner, A., Hartwigsen, G., & Roman Siebner, H. (2013). Right-hemispheric processing of non-linguistic word features: implications for mapping language recovery after stroke. Hum Brain Mapp, 34(6), 1293–1305.

Behrens, S. J. (1988). The role of the right hemisphere in the production of linguistic stress. Brain Lang, 33(1), 104–127.

Behrens, S. J. (1989). Characterizing sentence intonation in a right hemisphere-damaged population. Brain and language.

Borenstein, M., Hedges, L. V., Higgins, J. P. T., & Rothstein, H. R. (2011). Introduction to meta-analysis. John Wiley & Sons.

Brådvik, B., Dravins, C., Holtås, S., Rosen, I., Ryding, E., & Ingvar, D. H. (1991). Disturbances of speech prosody following right hemisphere infarcts. Acta Neurologica Scandinavica, 84(2), 114–126.

Bürkner, P.-C. (2017a). Advanced Bayesian Multilevel Modeling with the R Package brms. arXiv preprint arXiv:1705.11123.

Bürkner, P.-C. (2017b). brms: An R package for Bayesian multilevel models using Stan. Journal of Statistical Software, 80(1), 1–28.

Bürkner, P.-C., & Vuorre, M. (2018). Ordinal Regression Models in Psychology: A Tutorial. Advances in Methods and Practices in Psychological Science, 2515245918823199.

Cochran, W. G. (1954). The combination of estimates from different experiments. Biometrics, 10(1), 101–129.

Cohen, J. (1988). Statistical power analysis for the behavioral sciences. 2nd.

Cook, R. D., & Weisberg, S. (1982). Residuals and Influence in Regression. Chapman & Hall/CRC.

Cumming, G. (2014). The new statistics: why and how. Psychol Sci, 25(1), 7–29.

Cummins, N., Scherer, S., Krajewski, J., Schnieder, S., Epps, J., & Quatieri, T. F. (2015). A review of depression and suicide risk assessment using speech analysis. Speech Communication, 71, 10–49.

Degottex, G., Kane, J., Drugman, T., Raitio, T., & Scherer, S. (2014). Covarep?a collaborative voice analysis repository for speech technologies.

Egger, M., Smith, G. D., Schneider, M., & Minder, C. (1997). Bias in meta-analysis detected by a simple, graphical test. Bmj, 315(7109), 629–634.

Emmorey, K. D. (1987). The neurological substrates for prosodic aspects of speech. Brain and Language, 30(2), 305–320.

Erdfelder, E., Faul, F., & Buchner, A. (1996). GPOWER: A general power analysis program. Behavior research methods, instruments, & computers, 28(1), 1–11.

Forbes-Riley, K., & Litman, D. J. (2004). Predicting Emotion in Spoken Dialogue from Multiple Knowledge Sources.

Fusaroli, R., Lambrechts, A., Bang, D., Bowler, D. M., & Gaigg, S. B. (2017). Is voice a marker for Autism spectrum disorder? A systematic review and meta-analysis”. Autism Res, 10(3), 384–407.

Gandour, J., & Baum, S. R. (2001). Production of stress retraction by left- and right-hemisphere-damaged patients. Brain Lang, 79(3), 482–494.

Gandour, J., Larsen, J., Dechongkit, S., Ponglorpisit, S., & Khunadorn, F. (1995). Speech prosody in affective contexts in Thai patients with right hemisphere lesions. Brain and Language, 51(3), 422–443.

Gardner, H. (1994). The stories of the right hemisphere. Integrative views of motivation, cognition and emotion, 41, 57–69.

Gelman, A., Jakulin, A., Pittau, M. G., & Su, Y.-S. (2008). A weakly informative default prior distribution for logistic and other regression models. The Annals of Applied Statistics, 2(4), 1360–1383.

Gorelick, P. B., & Ross, E. D. (1987). The aprosodias: further functional-anatomical evidence for the organisation of affective language in the right hemisphere. Journal of Neurology, Neurosurgery & Psychiatry, 50(5), 553–560.

Guranski, K., & Podemski, R. (2015). Emotional prosody expression in acoustic analysis in patients with right hemisphere ischemic stroke. Neurologia i Neurochirurgia Polska, 49(2), 113–120.

Hedges, L. V., Tipton, E., & Johnson, M. C. (2010). Robust variance estimation in meta-regression with dependent effect size estimates. Res Synth Methods, 1, 39–65.

Hird, K., & Kirsner, K. (2003). The effect of right cerebral hemisphere damage on collaborative planning in conversation: an analysis of intentional structure. Clinical Linguistics & Phonetics, 17(4-5), 309–315.

Hoekert, M., Kahn, R. S., Pijnenborg, M., & Aleman, A. (2007). Impaired recognition and expression of emotional prosody in schizophrenia: review and meta-analysis. Schizophr Res, 96(1-3), 135–145.

Hughlings Jackson, J. (1879). On affections of speech from disease of the brain. Brain, 2, 203–222.

Jeffreys, H. (1998). The Theory of Probability. OUP Oxford.

Jiam, N. T., Caldwell, M., Deroche, M. L., Chatterjee, M., & Limb, C. J. (2017). Voice emotion perception and production in cochlear implant users. Hear Res, 352, 30–39.

Kass, R. E., & Raftery, A. E. (1995). Bayes factors. Journal of the American Statistical Association, 90(430), 773–795.

Kent, R. D., & Rosenbek, J. C. (1982). Prosodic disturbance and neurologic lesion. Brain and language.

Kotz, S. A., Meyer, M., Alter, K., Besson, M., von Cramon, D. Y., & Friederici, A. D. (2003). On the lateralization of emotional prosody: An event-related functional MR investigation. Brain and Language, 86(3), 366–376.

Leung, J. H., Purdy, S. C., Tippett, L. J., & Leão, S. H. (2017). Affective speech prosody perception and production in stroke patients with left-hemispheric damage and healthy controls. Brain Lang, 166, 19–28.

Liscombe, J., Venditti, J., & Hirschberg, J. B. (2003). Classifying subject ratings of emotional speech using acoustic features.

Macwhinney, B., Fromm, D., Forbes, M., & Holland, A. (2011). AphasiaBank: Methods for Studying Discourse. Aphasiology, 25(11), 1286–1307.

MacWhinney, B., Fromm, D., Rose, Y., & Bernstein Ratner, N. (2018). Fostering human rights through TalkBank. Int J Speech Lang Pathol, 20(1), 115–119.

Mahalanobis, P. C. (1936). On the generalized distance in statistics.

McCann, J., & Peppé, S. (2003). Prosody in autism spectrum disorders: a critical review. Int J Lang Commun Disord, 38(4), 325–350.

Monrad-Krohn, G. H. (1957). The Third Element of Speech: Prosody in the Neuro-Psychiatric Clinic. Journal of Mental Science, 103(431), 326–331.

Monrad-Krohn, G. H. (1947). The Prosodic Quality Of Speech And Its Disorders:(A Brief Survey From A Neurologist’s Point Of View). Acta Psychiatrica Scandinavica, 22(3-4), 255–269.

Moriarty, P. M., Vigeant, M., Liu, P., Gilmore, R., & Cole, P. (2018). Comparing theory, consensus, and perception to the acoustics of emotional speech. The Journal of the Acoustical Society of America, 144(3), 1841–1841.

Myers, P. S. (1999). Right hemisphere damage: Disorders of communication and cognition. Singular Pub.

Ouellette, G. P., & Baum, S. R. (1994). Acoustic analysis of prosodic cues in left- and right-hemisphere-damaged patients. Aphasiology, 8(3), 257–283.

Parola, A., Simonsen, A., Bliksted, V., & Fusaroli, R. (2019). Voice patterns in schizophrenia: A systematic review and Bayesian meta-analysis. bioRxiv, 583815.

Patel, S., Oishi, K., Wright, A., Sutherland-Foggio, H., Saxena, S., Sheppard, S. M. et al. (2018). Right Hemisphere Regions Critical for Expression of Emotion Through Prosody. Frontiers in Neurology, 9.

Pell, M. D. (1999a). Fundamental frequency encoding of linguistic and emotional prosody by right hemisphere-damaged speakers. Brain and Language, 69(2), 161–192.

Pell, M. D. (1999b). The temporal organization of affective and non-affective speech in patients with right-hemisphere infarcts. Cortex.

Peppé, S. J. E. (2009). Why is prosody in speech-language pathology so difficult. Int J Speech Lang Pathol, 11(4), 258–271.

Team, R. C. (2018). R: A language and environment for statistical computing. Vienna: R Foundation for Statistical Computing.

Rohatgi, A. (2014). Web Plot Digitizer. http.arohatgi.info/WebPlotDigitizer/app/(accessed June, 2.

Ross, E. D., & Monnot, M. (2008). Neurology of affective prosody and its functional-anatomic organization in right hemisphere. Brain Lang, 104(1), 51–74.

Ross, E. D., Harney, J. H., & Purdy, P. D. (1981). How the brain integrates affective and propositional language into a unified behavioral function: Hypothesis based on clinicoanatomic evidence. Archives of Neurology, 38(12), 745–748.

Ross, E. D., & Mesulam, M.-M. (1979). Dominant language functions of the right hemisphere?: Prosody and emotional gesturing. Archives of neurology, 36(3), 144–148.

Ross, E. D., Thompson, R. D., & Yenkosky, J. (1997). Lateralization of affective prosody in brain and the callosal integration of hemispheric language functions. Brain and language, 56(1), 27–54.

Ryalls, J., Joanette, Y., & Feldman, L. (1987). An acoustic comparison of normal and right-hemisphere-damaged speech prosody. Cortex, 23(4), 685–694.

Schuller, B., Steidl, S., Batliner, A., Vinciarelli, A., Scherer, K., Ringeval, F. et al. (2013). The INTERSPEECH 2013 computational paralinguistics challenge: social signals, conflict, emotion, autism.

Stoop, T. B., Moriarty, P., Vigeant, M., Gilmore, R., Liu, P., & Cole, P. (2018). Children’s ratings of vocal emotion intensity depend on the emotion spoken and speaker familiarity but not acoustic parameters. The Journal of the Acoustical Society of America, 144(3), 1965–1966.

Tompkins, C. A. (1995). Right hemisphere communication disorders: Theory and management. Singular Publishing Group.

Tsanas, A., Little, M. A., McSharry, P. E., & Ramig, L. O. (2010). Accurate telemonitoring of Parkinson’s disease progression by noninvasive speech tests. IEEE Trans Biomed Eng, 57(4), 884–893.

Vehtari, A., Gelman, A., & Gabry, J. (2017). Practical Bayesian model evaluation using leave-one-out cross-validation and WAIC. Statistics and Computing, 27(5), 1413–1432.

Viechtbauer, W. (2010). Conducting meta-analyses in R with the metafor package. J Stat Softw, 36(3), 1–48.

Wetzels, R., & Wagenmakers, E. J. (2012). A default Bayesian hypothesis test for correlations and partial correlations. Psychon Bull Rev, 19(6), 1057–1064.

Williams, D. R., Rast, P., & Bürkner, P.-C. (2018). Bayesian Meta-Analysis with Weakly Informative Prior Distributions.

Winner, E., Brownell, H., Happé, F., Blum, A., & Pincus, D. (1998). Distinguishing lies from jokes: Theory of mind deficits and discourse interpretation in right hemisphere brain-damaged patients. Brain and language, 62(1), 89–106.

Wright, A., Saxena, S., Sheppard, S. M., & Hillis, A. E. (2018). Selective impairments in components of affective prosody in neurologically impaired individuals. Brain and Cognition, 124, 29–36.

Yang, S. Y., & Van Lancker Sidtis, D. (2016). Production of Korean Idiomatic Utterances Following Left- and Right-Hemisphere Damage: Acoustic Studies. J Speech Lang Hear Res, 59(2), 267–280.

Yao, Y., Vehtari, A., Simpson, D., & Gelman, A. (2018). Using stacking to average Bayesian predictive distributions. Bayesian Analysis.

